# CXCL13 levels in cerebrospinal fluids of patients with multiple sclerosis: The role of Borrelia proteins in neuro infections

**DOI:** 10.1101/2023.01.05.522966

**Authors:** Şeyda Karabörk, Şule Aydin Türkoğlu, Serpil Yildiz, Fatma Sirmatel

## Abstract

In the present study, the purpose was to study anti-Borrelia antibodies with both ELISA and WB for the control of Lyme Disease in CSF samples obtained from patients diagnosed with MS, and to determine the relationship between them by investigating the CXCL13 levels. A total of 43 CSF samples taken from patients diagnosed with MS and PTS. The obtained data were statistically analyzed with the Spearman Rank Correlation Test and p<0.05 value was considered statistically significant. Especially 5 antigens (p19, p20, p21, p58, and OspC) were found to be positive as a result of the confirmation of the samples that were positive for Borrelia antibodies with the WB method. When the results of the study were evaluated, the Borrelia antibodies that were found positive by ELISA and high CXCL13 in CSF samples of MS patients proved once again that CXCL13 is still the best biomarker for LNB. The presence of Borrelia antibodies, which were found to be positive with the ELISA test in CSF samples of MS patients, was confirmed with WB. The coexistence of high CXCL13 levels in the same samples suggests that LNB may also play a role in the etiopathogenesis of MS and CXCL13 may be a potential biomarker in this respect. Also, with the positive detection of OspC and p58 WB bands in the majority of cases, we think that these two protein bands will shed light on borrelia studies in CSF in neurodegenerative diseases and can be used as a potential biomarker in diagnosis.

## INTRODUCTION

Multiple Sclerosis (MS) is a heterogeneous, long-term, autoimmune, neurodegenerative, demyelinating disease of the Central Nervous System (CNS) most common in young adults, which can manifest itself with chronic, inflammatory, episodic neurological deficits and a neuroanatomical localization (1, 2). Many environmental and genetic factors are considered to play roles in its pathogenesis (3). Relapsing-Remitting MS (RRMS) is the form that progresses with attacks and remissions in the most common form of the disease. The cases without typical MS findings and normal clinical findings, but with MS-like demyelinating lesions on cranial MRI imaging performed for any reason and other causes excluded, have been defined as a Radiologically Isolated Syndrome (RIS) in recent years (4). As damage to brain tissue accumulates over the years, patients often develop Secondary Progressive MS (SPMS), whose neurological deficits worsen over time. Two different pathological processes were identified in patients with RRMS (inflammation/demyelination, which is dominant, and axonal degeneration, which is considered to be responsible for SPMS). The diagnostic criteria of MS are based on the demonstration of lesions in different CNS and clinical evaluations (5, 6, 7). In recent studies, many infectious and non-infectious environmental factors (e.g., EBV, vitamin D, smoking, and toxins) were blamed for the formation of MS (8, 9, 10, 11). There is a female predominance in MS and the young population is particularly affected. In the differential diagnosis of MS, there are rheumatological causes such as systemic lupus (SLE), APS, Behçet’s Disease, Sjögren’s Syndrome, isolated CNS vasculitis with similar clinical and MRI findings, and infectious causes such as neurosarcoidosis, neuroborreliosis, and syphilis (12–14). For this reason, when diagnosing MS, it is necessary to investigate other MS-like clinical and MRI-like causes. Especially, examination of infectious agents and autoimmune and immunological markers is important in this regard. In previous studies that investigate MS activity, cytokines and chemokines, especially in the cerebrospinal fluid (CSF), were studied as potential markers for MS. These cytokines and chemokines are important mediators involved in the immune response as modulators of cell recruitment and migration to sites of inflammation. An increase in pro-inflammatory is seen during MS exacerbations and anti-inflammatory cytokines and chemokines are decreased. Recent studies show that the levels of some chemokines and chemokine receptors are increased in the blood and CSF of MS patients (15, 16). Known as chemokine ligand 13, CXCL13 is a B-cell chemoattractant expressed in structural lymphoid organs and promotes the formation of B-cell follicles, which is important in the pathophysiology of MS. CXCL13 is elevated in the CSF of patients with various forms of MS and its levels appear to correlate with disease activity (17, 18). The detection of CXCL13 in CSF is a promising candidate to be a CSF biomarker for the evaluation of MS activity. However, studies conducted on this subject are insufficient and the results are not yet reliable enough to recommend its use in routine clinical practice. However, recent studies are showing that CXCL13 can be used as a marker even in the disease course of MS (17, 19, 20).

Lyme borreliosis is the most common vector-borne infectious disease in Europe and North America, transmitted by the contact and/or bite of ticks infected with the spirochete bacterium *Borrelia burgdorferi (B. burgdorferi)* (21, 22). In untreated or inadequately treated cases, Lyme Disease can cause numerous systemic clinical manifestations (23). Lyme Neuroborreliosis (LNB) is a common disease in Europe as well as in Sweden. It is already known that *B. burgdorferi*, which creates an innate immune response by interacting with the recognition receptors called Toll-Like Receptor 2 (TLR2), plays roles in the pathogenesis of LNB causing B and T cells to be activated in the CNS and recruited to these cells (24, 25). Since CXCL13 is an important molecule in B cell recruitment to the CNS, previous studies showed that it is found in high CXCL13 concentrations in the CSF of both children and adults with LNB. There are studies in which the CXCL13 level is elevated in CSF before detecting specific antibodies to *B.burgdorferi* produced intrathecally. For this reason, CXCL13 is accepted as a diagnostic marker in acute LNB (26, 27).

The similarity of the immune mechanism/immune pathogenesis of MS and Lyme Disease, and the fact that both are neurological diseases, make the diagnosis difficult in some cases. Especially the fact that Lyme Disease can mimic many diseases and sometimes show common symptoms that can cause certain problems. The high prevalence of both MS and Lyme Diseases in the city of Bolu, and the high number of patients with MS and Lyme (28, 29), it is necessary and important to focus on this issue and to use common or supportive biomarkers for rapid diagnosis of these diseases.

Pseudotumor Cerebri (PTS) is a disease of which the pathophysiology remains unclear. This neurodegenerative disorder is characterized by impaired CSF absorption and inflammatory changes (30). Although it is clinically characterized by headache, papilledema, vision loss, and pulsatile tinnitus, no explanatory cause can be detected in radiological imaging and CSF examination (31). Many studies were conducted on cytokines and chemokines as biomarker candidates in the progression, recovery, and even short-term diagnosis of neurodegenerative diseases such as MS and PTS (32). However, chemokine studies in which these two diseases exist together and especially CSF are examined are very few. For this reason, in the present study, the purpose was to study Anti-Borrelia tests for the control of Lyme Disease in CSF samples of patients diagnosed with MS and PTS for 4 years in the Neurology Clinic of our hospital and to screen for CXCL13, which is used in the diagnosis of both (especially Neuroborreliosis). It is considered that CXCL13 CSF associated with LNB can be determined, especially in neuroinflammatory and neurodegenerative diseases such as MS by evaluating CXCL13 levels in the CSF of MS and PTS patients, and a new diagnosis or treatment strategy can be developed that clinicians can use in the future.

## MATERIAL & METHOD

The approval for the study was obtained with the number 2018/100 from the Regional Ethics Committee. The study included MS, MS-like cases (n=23, n=5, respectively) taken by Lumbar puncture (LP) from the patients who applied to Bolu Abant İzzet Baysal University, Training and Research Hospital, Neurology Clinic between 2013 and 2017 for MS differential diagnosis. A total of 43 cases were included in the study, including CSF samples of cases diagnosed as PTS (n=15) and performed with evacuation LP and stored at −80°C. The patients were classified as MS-diagnosed and non-MS (RIS) according to McDonald Criteria. The socio-demographic characteristics of the patients were obtained from the archives as CSF biochemistry values (Albumin, Glucose, LDH, protein), and serum IgG and IgM values measured by the *B. burgdorferi* ELISA method were recorded retrospectively and analyzed to add strength to the study. The correlation was provided with the EDSS scores of MS patients. All CSF samples taken until the time of the study were stored at −80°C under appropriate conditions. All tests of the study such as CXCL13 ELISA, CSF Borrelia IgG, IgM ELISA, and CSF Borrelia IgG and IgM Western blot (Euroimmun, Lubeck, Germany) were conducted in line with company recommendations and the results were evaluated and recorded. For the CXCL13 test, <20 pg/ml was considered normal, ≥20 bis <30 pg/ml borderline/grey zone, ≥30 bis ≤100 pg/ml increasing/rising “Increased”; and >100 pg/ml values were considered “Strongly increased” showing acute neuroborreliosis.

### For Borrelia IgG and Borrelia IgM Elisa test

a test kit that contained *B. burgdorferi, B. afzelii, B. garinii,* Outer surface protein C (OspC), and Borrelia Variable Surface Lipoprotein E (VlsE) antigens was preferred. The samples determined as positive or borderline according to the Elisa Test results were confirmed with the Western Blot (WB) Test, considering the algorithm called the “two-tier” strategy in the diagnosis of Borrelia according to the Centers for Disease Control and Prevention (CDC) data. According to the immunoblot test method, the antigens (p18, p19, p20, p21, p58, p39, p83) of four different Borrelia strains (*B. burgdorferi, B. afzelii, B. garinii, B. spielmanii)* that are pathogenic in humans and outer surface protein, which is major in serology lipoproteins called OspC and VlsE were evaluated. The results were noted and compared according to the methods.

### Statistical analysis

For statistical analysis, the results were given as mean ± SD or median (min-max) according to their normal distribution. All statistical analyzes were performed by using the “SPSS for Windows 17.0” (SPSS, Chicago, IL, USA) program. The Student-***t*** test was used to compare the age distributions between MS and RIS groups. The Pearson Chi-Square Test was used for gender comparison. The distribution of normality continuous variables was determined with the Shapiro-Wilk Test. The analysis of CXCL13 concentrations, *B. burgdorferi* positivity, clinical parameters, and CSF laboratory abnormalities was performed by using the Spearman Rank Correlation Test, and p<0.05 was considered statistically significant.

## RESULTS

The MS group (11 females, 12 males) and the RIS group (2 females, 3 males) were classified as demyelinating groups. The demographic data of the patients are summarized in Table 1. A statistically significant difference was detected between the mean age of the demyelinating group (13 females, 15 males) and the PTS group (10 females, 5 males), respectively, with a mean (±SD) 35 (±12), 45 (±11) (p=0.01). No difference was detected between the two groups in terms of gender distribution (p=0.2). There were no cells in the CSF in the CSF examinations of all cases. No statistically significant differences were detected in CSF biochemistry values between the groups (Table 2). The serum *B. burgdorferi* results of all cases that were recorded in the system and measured with the ELISA Method were recorded. Although serum *B. burgdorferi* IgM was negative in all cases, serum *B. burgdorferi* IgG was positive in 5 MS, 1 RIS and 1 PTS cases. Although *B. burgdorferi* IgM studied in the CSF of the MS and RIS group was found to be negative in all cases, *B. burgdorferi* IgG was detected as positive in 5 of the MS group. When the CXCL13 values were evaluated, the cases with higher than 100 pg/ml were encountered with highly suspicious Lyme. When 3 cases were examined, it was found that 2 of them had *B.burgdorferi* IgG positive in serum, which was measured by ELISA, and *B. burgdorferi* IgG positive was found in the CSF of these cases as a result of WB verification. In the WB band examination of these cases, p19, p20, p21, p58, and OspC bands were positive. In one case, although *B. burgdorferi* IgG measured with serum ELISA was positive, both *B. burgdorferi* IgG and IgM were negative when WB confirmation was performed in the CSF sample. The presence of p20, p58, and OspC bands was detected in the WB band examination of this case. Also, when 2 cases with moderately suspicious Lyme with CXCL13 values above 30-100 pg/ml were examined, although *B. burgdorferi* IgG, which was measured by ELISA, was positive in the serum, WB confirmation was performed in the CSF. As a result, *B. burgdorferi* IgG and IgM were negative in the WB test. P58 and OspC bands were also observed in the WB band examination of these cases (Table 3). CSF CXCL13 values of all three groups were 8 (2-125), 7 (3-17), and 2 (1-11), respectively, as median (min-max) and no significant differences were detected between MS and RIS groups (p=0.6). A statistically significant difference was detected between the demyelinating group and PTS (p=0.001), and the CXCL13 value was significantly higher in the demyelinating group than in the PTS group (Table 4). When the correlation between CXCL13 and CSF and serum *B. burgdorferi* positivity of the cases was examined, it was found that there was a mild-moderate strong positive correlation with *B. burgdorferi* IgG positivity in CSF, which was measured with both ELISA and WB verification, respectively (p<0.001, p. =0.02) (Rho=0.6, Rho=0.5) (Table 5).

**Table 1.** The demographic data and clinical characteristics of the patients

| Groups<br>(n=43) | Demyelinating (n=28) |  | Demyelinating<br>(n=28) | PTS<br>n=15 |
| --- | --- | --- | --- | --- |
|  | MS (n=23) | RIS (n=5) |  |  |
| Mean Age ± SD | 33 ± 11 | 43 ± 16 | 35 ± 12 | 45 ± 11 |
|  | p=0.1 |  | p=0.01<sup>±</sup> |  |
| Female | 11 | 2 | 13 | 10 |
| Male | 12 | 3 | 15 | 5 |
|  | p=0.8 |  | p=0.2* |  |

MS: Multiple Sclerosis, RIS: Radiological Isolated Syndrome, <sup>±</sup>T Tests, \*Pearson Chi-Square

**Table 2.** The comparison of CSF biochemistry values between groups

|  | BOS<br>Protein | Serum<br>Protein | BOS<br>Albumin | Serum<br>Albumin | BOS<br>Glucose | Serum<br>Glucose | BOS<br>LDH | BOS<br>Chlorine |
| --- | --- | --- | --- | --- | --- | --- | --- | --- |
| Case | 30<br>(15-226) | 7<br>(6-71) | 15<br>(1-30) | 4<br>(4-10) | 71<br>(50-126) | 104<br>(98-110) | 16<br>(10-41) | 125<br>(119-131) |
| PTS | 36<br>(17-174) | 7<br>(7-8) | 16<br>(1-30) | 4<br>(4-5) | 63<br>(54-83) | 107<br>(104-112) | 18<br>(10-31) | 124<br>(120-130) |
| p±= | 0.7 | 0.9 | 0.5 | 0.1 | 0.3 | 0.07 | 0.5 | 1 |

The Case: Demyelinating Group, PTS: Pseudotumor Cerebri ± Mann-Whitney-U

**Table 3.** CXCL13, CSF Borrelia ELISA, and WB values of the case and control groups

| The<br>Case | EDSS | CXCL13<br>(pg/ml) | Lyme<br>IgG | Lyme<br>IgM | WB IgG | WB<br>IgM | IgG WB | OCB<br>Type | Index |
| --- | --- | --- | --- | --- | --- | --- | --- | --- | --- |
| 1RRMS | 2 | 14.5 | Negative | Negative | Negative | Negative | None | Type<br>2 | 1.17 |
| 2RRMS | 2 | 11 | Negative | Negative | Negative | Negative | p58 | Type<br>1 | 0.66 |
| 3RRMS | 1 | 8.2 | Negative | Negative | Negative | Negative | OspC | Type<br>2 | 1.17 |
| 4RRMS | 1 | 12 | Negative | Negative | Negative | Negative | OspC | Type<br>2 | 0.73 |
| 5RRMS | 3 | 7 | Negative | Negative | Negative | Negative | p58 and<br>OspC | Type<br>2 | 0.67 |
| 6RRMS | 1 | 92 | Positive | Negative | Negative | Negative | p58 and<br>OspC | Type<br>2 | 1.67 |
| 7RRMS | 3 | 125 | Positive | Negative | Positive | Negative | p19. p20.<br>p21. p58<br>and OspC | Type<br>2 | 0.68 |
| 8RRMS | 1 | 111 | Positive | Negative | Positive | Negative | p19. p20.<br>p21. p58<br>and OspC | Type<br>3 | 1.16 |
| 9RRMS | 2 | 2.2 | Negative | Negative | Negative | Negative | p19. p20.<br>p21 and<br>p58 | Type<br>2 | 0.94 |
| 10RRMS | 1 | 4 | Negative | Negative | Negative | Negative | p19. p20.<br>p58 and<br>OspC | Type<br>1 | 0.53 |
| 11RRMS | 1 | 31.2 | Positive | Negative | Negative | Negative | p58 and<br>OspC | Type<br>2 | 2 |
| 12RRMS | 2 | 20 | Negative | Negative | Negative | Negative | p58 and<br>OspC | Type<br>1 | 0.72 |
| 13RRMS | 3.5 | 108 | Positive | Negative | Negative | Negative | p20. p58<br>and OspC | Type<br>1 | 0.57 |
| 14RRMS | 1 | 10 | Negative | Negative | Negative | Negative | p20. p58<br>and OspC | Type<br>4 | 0.58 |
| 15RRMS | 1 | 4.6 | Negative | Negative | Negative | Negative | p58 and<br>OspC | Type<br>1 | 0.55 |
| 16RRMS | 2 | 1.8 | Negative | Negative | Negative | Negative | p58 and<br>OspC | Type<br>4 | 0.6 |
| 17RRMS | 1 | 5 | Negative | Negative | Negative | Negative | p58 and<br>OspC | Type<br>2 | 1.16 |
| 18RRMS | 1 | 7 | Negative | Negative | Negative | Negative | p58 and<br>OspC | Type<br>2 | 1.12 |
| 19RRMS | 1 | 18 | Negative | Negative | Negative | Negative | p58 and<br>OspC | Type<br>2 | 0.94 |
| 20RRMS | 1 | 2 | Negative | Negative | Negative | Negative | p58 and<br>OspC | Type<br>2 | 1.36 |
| 21RRMS | 2 | 1.5 | Negative | Negative | Negative | Negative | p58 and<br>OspC | Type<br>2 | 1.14 |
| 22RRMS | 2 | 2.8 | Negative | Negative | Negative | Negative | p58 and<br>OspC | Type<br>2 | 1.1 |
| 23RRMS | 1 | 8 | Negative | Negative | Negative | Negative | p58.<br>OspC.<br>p39 and<br>p41 | Type<br>2 | 0.98 |
| 24RIS | 0 | 17 | Negative | Negative | Negative | Negative | p58 and<br>OspC |  |  |
| 25RIS | 0 | 13 | Negative | Negative | Negative | Negative | p58 and<br>OspC |  |  |
| 26RIS | 0 | 2.9 | Negative | Negative | Negative | Negative | p58 and<br>OspC |  |  |
| 27RIS | 0 | 4.5 | Negative | Negative | Negative | Negative | p58 and<br>OspC |  |  |
| 28RIS | 0 | 7 | Negative | Negative | Negative | Negative | p58 and<br>OspC |  |  |
| 29PTS |  | 1.94 | Negative | Negative |  |  |  |  |  |
| 30PTS |  | 1.74 | Negative | Negative |  |  |  |  |  |
| 31PTS |  | 2.22 | Negative | Negative |  |  |  |  |  |
| 32PTS |  | 1.21 | Negative | Negative |  |  |  |  |  |
| 33PTS |  | 1.15 | Negative | Negative |  |  |  |  |  |
| 34PTS |  | 3.24 | Negative | Negative |  |  |  |  |  |
| 35PTS |  | 4.02 | Negative | Negative |  |  |  |  |  |
| 36PTS |  | 1.26 | Negative | Negative |  |  |  |  |  |
| 37PTS |  | 2.29 | Negative | Negative |  |  |  |  |  |
| 38PTS |  | 11 | Negative | Negative |  |  |  |  |  |
| 39PTS |  | 4.5 | Negative | Negative |  |  |  |  |  |
| 40PTS |  | 8.1 | Negative | Negative |  |  |  |  |  |
| 41PTS |  | 1.98 | Negative | Negative |  |  |  |  |  |
| 42PTS |  | 4.22 | Negative | Negative |  |  |  |  |  |
| 43PTS |  | 2.23 | Negative | Negative |  |  |  |  |  |

RRMS: Relapsing-Remitting Multiple Sclerosis, RIS: Radiologically Isolated Syndrome, OCB: Oligoclonal band

**Table 4.** The intergroup comparison of CSF CXCL13 values

| Group n=43 | Demyelinating group n=28 |  | PTS n=15 |
| --- | --- | --- | --- |
| CXCL13 (pg/ml)<br>median (min-max) | MS | RIS | 2 (1-11) |
|  | 8 (2-125) | 7 (3-17), |  |
|  | p=0.6° |  |  |
|  | 8 (2-125) |  |  |
|  | p=0.001* |  |  |

Demyelinating Group, MS: Multiple Sclerosis, RIS: Radiologically Isolated Syndrome, PTS: Pseudotumor Cerebri, ° Kolmogorov-Smirnov Z \*Mann-Whitney U

**Table 5.**
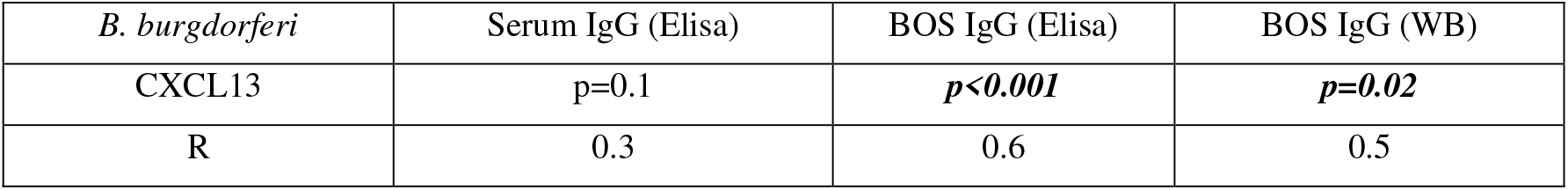
The relationship between CSF CXCL13 and *B. burgdorferi* positivity

| *B. burgdorferi* | Serum IgG (Elisa) | BOS IgG (Elisa) | BOS IgG (WB) |
| --- | --- | --- | --- |
| CXCL13 | p=0.1 | **p<0.001** | **p=0.02** |
| R | 0.3 | 0.6 | 0.5 |

## DISCUSSION

Chemokines are a family of small molecular weight proteins that are responsible for the migration of cells in the immune response. They are known for their ability to act as chemoattractants. As with lymphoproliferative diseases, many diseases (including infectious diseases) chronic inflammation, and autoimmune diseases, were associated with abnormal chemokine and cytokine production. CXCL13 and its receptor and the G-protein coupled receptor (GPCR) CXCR5 are well known to play essential roles in inflammatory, infectious, and immune responses (17, 33). The CXCL13:CXCR5 axis plays roles in both physiological and pathological humoral immunity. For this reason, it is involved in the pathogenesis of infectious diseases. Especially among these infectious diseases, the most emphasized is the role of CXCL13 in the Human Immunodeficiency Virus in the early and chronic period, even in children. As a result of previous studies, CXCL13 was identified as a strong biomarker candidate in line with the literature data (33, 34). Similar contributions of CXCL13 to the formation of lymphoid structures in the immune system were also demonstrated in Sjögren’s syndrome, autoimmune thyroiditis, myasthenia gravis, systemic lupus erythematosus (SLE), and MS. As a result, CXCL13 expression was proposed as a biomarker for MS disease exacerbation and diagnosis and progression in such conditions (33). Considering the central role it plays in humoral immunity, it was observed that CXCL13:CXCR5 was studied especially on the stages of Syphilis and Lyme Disease (Lyme neuroborreliosis and neurosyphilis), which affect the CNS. It was shown that CXCL13 is overexpressed in the muscles of monkeys chronically infected with *B. burgdorferi* and subsequently contributes to the formation of ectopic germinal centers in the CNS. Interestingly, although infection with *B. burgdorferi* did not seem to have any impacts on plasma CXCL13 levels, CXCL13 levels in CSF were found to be 100-fold higher than in plasma when CNS infection with bacteria was established (27, 35). In a meta-analysis study that was conducted after this determination, the CSF CXCL13 level was proposed as a diagnostic biomarker for neuroborreliosis by calculating the sensitivity and specificity of 89% and 96% for CXCL13, which were reviewed in 18 studies (27). CXCL13 levels were also found to be elevated in CSF samples of MS patients (27, 36). In our study, as in the literature, the CXCL13 level was found to be statistically significantly higher in the demyelinating group in CSF than in the PTS group (p=0.001). The detection of CXCL13 in CSF suggests that this chemokine also plays roles as a possible marker of disease severity. It was also shown that CSF CXCL13 levels are associated with the course of MS disease (18, 37). These results suggest that CSF CXCL13 must be used in clinical practice to diagnose MS and CXCL13 has the potential to be a biomarker for response to drug therapy and the progression of MS disease. However, further studies are required because of the limited number of studies in the literature to confirm this. Another aspect of our study was that the screening of Borrelia antibodies together with CXCL13 in cases with MS, RIS, and PTS and their confirmation with the WB test brought a novelty in this respect. It is noteworthy that positivity/positive bands are observed in some of the specific protein antigens included in the WB test. Each of the p19, p20, p21, p39, p41, p58, and OspC bands in our cases had different characteristics. As it is already known, antibodies against OspC encoded on plasmids are the most important serological marker to detect Borrelia infections (sensitivity >90%). Because OspC is one of the major lipoproteins expressed on the surface of *B. burgdorferi* during the early phase of cases after tick exposure. OspC is required for *B. burgdorferi* to cause infection and is one of the important criteria used in the diagnosis (38). Although 5 out of 5 bands were not positive in MS and RIS individuals who constituted the experimental group of the present study, OspC was found positive in 26 of 28 cases. This shows that they met Borrelia at some point in their lives. It is already known that p58, which is a chromosomal protein from the integral outer proteins, is highly specific for Borrelia, but functions only with oligopeptide proteins whose function has not been clarified yet, and it is also found in *B.garinii* species other than *B.burgdorferi,* and this species is also neurotropic (38). In our cases, when the MS and RIS groups were considered, p58 positive band was detected in the cases with negative or positive WB results. For this reason, a specific interpretation could not be reached for the p58 protein band. Nothing can be said about the early or late stage of Lyme Disease. It is seen that the specificity and function of the positive bands associated with p19 OspE are not known. It is already known that p21 from other bats binds a protein called decorin, which is responsible for binding Type I collagen and Tumor Growth Factor (TGF-ß) in the connective tissue on the host cell especially on the skin, which is an important characteristic for cases of Erythema migrants (EM), the first stage of Lyme Disease. P39, which is also known as Borrelia membrane protein A (BmpA), is one of the antigens that become positive a few weeks after tick bite/tick contact, which is known to be highly specific for Borrelia. Although the specificity of the flagellin protein p41 is not clear, it is the first antigen to be formed in the WB IgM test. It can also cross-react with flagellate/flagellate bacteria or other spirochetes (40). This suggests that the positivity of p41, which was detected in only 1 of our cases, may be because of another spirochete infection and/or the presence of a flagellated bacterial infection. The present study has led us to consider Borrelia infection in neurodegenerative diseases such as MS in the city of Bolu, which is a tick-endemic region. When the WB kits used in the study were evaluated according to the European İmmunoblot All Antigen Criteria, ≥ 2 bands for IgG and ≥ 1 band for IgM are considered positive (41). In the Recombinant Immunoblot Criterion of Goettner et al. and Schulte-Spechtel et al., ≥ 2 bands for IgG and ≥ 2 bands for IgM are evaluated as positive (42, 43).

Considering the European and recombinant-based immunoblot tests, almost all of our MS and RIS cases are considered positive. This once again highlights that Borrelia antibodies must be evaluated while investigating the underlying mechanisms in MS cases. Another point that makes us think is the necessity of preparing new diagnostic kits containing the strains for a special evaluation and diagnosis for our country, considering the CDC, Europe, and Recombinant tests. As a result, in the present study, the finding of high CXCL13 in CSF samples of MS patients with positive Borrelia ELISA tests once again proved that CXCL13 is still the best biomarker for LNB. It was pointed out that Borrelia antibodies must be investigated not only in serum samples but also in CSF samples if necessary, when diagnosing neuroborreliosis. The detection of CXCL13 in CSF also suggests a role for this chemokine as a possible marker of disease severity, which suggests that CSF CXCL13 must be used in clinical practice for the diagnosis of MS and that CXCL13 also has the potential to be a biomarker for response to drug therapy and progression of MS disease, but further studies are needed to confirm this. The most important limitation of this study was that it had a small sample. Prospective studies with more samples are needed in the future.

### Conclusion

When the results of the present study were evaluated, the finding of high CXCL13 in CSF samples of MS patients with positive Borrelia ELISA tests once again proved that CXCL13 is still the best marker/biomarker for LNB. It was pointed out that Borrelia antibodies must be investigated not only in serum samples but also in CSF samples if necessary when diagnosing neuroborreliosis. We believe that CSF CXCL13 measurement will help diagnostic and prognostic studies in MS, disease severity and course, and support future treatment options/decisions. We also think that these two protein bands will shed light on Borrelia studies in CSF with the positive detection of OspC and p58 WB bands in most cases. According to the results obtained here, we predict that OspC and p58 can be used as potential biomarkers in the diagnosis. There is a need for further studies with larger sample numbers.

This research is supported by Bolu Abant İzzet Baysal University Scientific Research Projects (No. 2017.08.32.1257) and we are very grateful in this regard.

## Source of funding

None

## Conflict of Interest

None to declare

## Author Contributions

Concept – S.A.T; S.K Design – S.K.; Supervision – S.Y; F.S. Materials Data Collection and/or Processing – S.A.T Statistics – S.A.T.; S.Y. Literature Search – S.K; S.A.T. S.Y.; Writing – S.K; S.A.T. S.Y Critical Reviews – S.Y; F:S

## Notes

Footnote: The present study was supported (2017.08.32.1257) in the scope of Scientific Research Projects at Bolu Abant İzzet Baysal University.

